# Sex-specific maturational trajectory of endocannabinoid plasticity in the rat prefrontal cortex

**DOI:** 10.1101/2020.10.09.332965

**Authors:** Axel Bernabeu, Anissa Bara, Antonia Manduca, Milene Borsoi, Olivier Lassalle, Anne-Laure Pelissier-Alicot, Olivier JJ Manzoni

## Abstract

The prefrontal cortex (PFC) develops until early adulthood in rodents and humans, but how synaptic plasticity evolves throughout postnatal development is not known. Here, we used a cross-sectional approach to establish the postnatal maturational trajectories of intrinsic properties and synaptic plasticity in the PFC of rats of both sexes. We found that while layer 5 PFC pyramidal neurons from rats of both sexes displayed similar current-voltage relationships, rheobases and resting potentials across all age groups, excitability was lower in female adults compared to the other developmental stages. NMDAR-dependent long-term potentiation and mGluR2/3-mediated long-term depression (LTD) were equally expressed at the juvenile, pubescent and adult developmental stages in animals of both sexes. However, the developmental course of endocannabinoid (eCB)-mediated LTD was sexually dimorphic. First, eCB-LTD emerged during the juvenile period in females. However, although CB1Rs were functional in both sexes at all developmental stages, eCB-LTD’s first emerged during pubescence in male. Second, eCB-LTD engaged distinct receptors in males and females depending on their developmental stages. Female rats employ both CB1R and TRPV1R to produce eCB-LTD at the juvenile stage but solely CB1R at pubescence followed by only TRPV1R at adulthood. In contrast, in pubescent and adult males eCB-LTD always and exclusively depended on CB1R. Pharmacological blockade of 2AG’s principal degrading enzyme allowed incompetent male juvenile synapses to express eCB-LTD. The data reveal different maturational trajectories in the PFC of male and female rats and provide new cellular substrates to the sex-specific behavioral and synaptic abnormalities caused by adolescent exposure to cannabinoids.

## Introduction

The endocannabinoid (eCB) system, comprised of the cannabinoid receptors (CB1R and CB2R) and the vanilloid receptor TRPV1R, as well as the proteins synthesizing, transporting and degrading the endogenous cannabinoids anandamide (AEA) and 2-arachidonoylglycerol (2-AG), contributes to development beginning at ontogenesis (Hurd et al. 2019) and continuing throughout the post-natal neurodevelopmental period (Harkany et al. 2007, 2008; Fride 2008; Jutras-Aswad et al. 2009; Galve-Roperh et al. 2013)(Lu and Mackie 2020)(Bossong and Niesink 2010). The eCB system is highly influenced by sex and hormonal regulation (Cooper and Craft, 2018; Krebs-Kraft et al., 2010). For example, in the CNS, circulating levels eCBs and the expression of their major receptor CB1R, vary with sex and during the estrous cycle (Rodríguez de Fonseca et al., 1994; González et al., 2000, Bradshaw et al., 2006). Not surprisingly, human and rodent studies consistently indicate sex differences in the short and long-term effects of cannabis and the trajectory of cannabis use (Stinson et al. 2006; Schneider 2008; Schepis et al. 2011; Craft et al. 2013; Lubman et al. 2015; Cuttler et al. 2016).

The prefrontal cortex (PFC) underlies multiple higher functions such as working memory, reasoning, cognitive flexibility and emotionally-guided behaviors (Goldman-Rakic 1991; Seamans et al. 1995), and continues to mature until the end of adolescence. Emerging evidence pinpoint adolescence as a sensitive period of development (Fuhrmann et al. 2015), a postnatal “period of heightened malleability’ (Steinberg and Morris 2001) during which external stimuli shape changes in brain structure and function. Illuminating the maturational sequence of the PFC is of particular significance in understanding how sensitive periods contribute to physiological development and the pathological processes that have their roots in early life adversity.

During adolescence, the maturation of corticolimbic areas implicates the endocannabinoid system, notably in the PFC (Meyer et al. 2018). The PFC is also a site of dense expression of CB1R (Marsicano and Lutz 1999) and is exquisitely sensitive to various synaptopathies which manifest as a variety of cognitive disease states (Scheyer et al. 2017). Exogenous cannabinoid exposure during the prenatal or the perinatal period results in significant abnormalities in PFC synaptic function which underlie long-lasting behavioral alterations at adulthood in cannabinoid-exposed rats (Manduca et al. n.d.; Bara et al. 2018; Scheyer et al. 2019, 2020a). In spite of substantial evidence that cannabinoid exposure during adolescence affects males and/or females (Borsoi et al. 2019)(Schneider 2008; Cass et al. 2014; Renard et al. 2016, 2016), the normal maturation profile of the endocannabinoid system in the PFC remains obscure.

Here, we applied a cross-sectional strategy to explore the maturational sequence of PFC layer 5 (L5) pyramidal neurons and their excitatory synapses in male and female rats. We report period and sex specific differences in the development and the receptors mediating endocannabinoid-mediated long-term depression (LTD) in the rat PFC. These differences were not apparent in the nucleus accumbens, a cognate structure of the mesocorticolimbic system. In contrast, at the same PFC excitatory synapses, neither long-term potentiation (LTP) nor type 2 mGluR-LTD varied during adolescent maturation in either sex. The intrinsic properties of L5 pyramidal cells also displayed a degree of sex-specific maturation while basic synaptic properties were invariant. By revealing sex differences in the maturational trajectories of the rat PFC, these data add new fundamental bases for the idea that adolescence is a sex-specific sensitive period and may provide novel substrates to the cellular and behavioral sex differences in the effect of adolescent cannabinoid exposure.

## Materials and Methods

Further information and requests for resources should be directed to the Lead Contact, Olivier J.J. Manzoni.

### Animals

Animals were treated in compliance with the European Communities Council Directive (86/609/EEC) and the United States NIH Guide for the Care and Use of Laboratory Animals. The French Ethical committee authorized the project “Exposition Périnatale aux cannabimimétiques” (APAFIS#18476-2019022510121076 v3). All rats were obtained from Janvier Labs and group-housed with 12h light/dark cycles with ad libitum access to food and water. All groups represent data from a minimum of 2 litters. Female data were collected blind to the estrous cycle. Female and male rats were classified based on the timing of their pubertal maturation. The pubertal period in female rats (approximately postnatal day (P) 28 to P 40) was determined by vaginal opening and first estrus. Balano preputial separation indicated pubertal onset in male rats (around P 40), and sexual maturity is indicated by the presence of mature spermatozoa in the vas deferens, which is achieved around P 60 (Schneider 2008). Thus, the female ages groups were Juvenile 21<P<28; Pubescent 30<P<50 and Adult at P >90. Male groups were: Juvenile 21<P<38; Pubescent 40<P<60 and Adult P >90.

### Slice preparation

Adult male and female rats were anesthetized with isoflurane and killed as previously described (Bara et al. 2018; Borsoi et al. 2019; Scheyer et al. 2019, 2020a, 2020b). The brain was sliced (300 µm) in the coronal plane with a vibratome (Integraslice, Campden Instruments) in a sucrose-based solution at 4°C (in mM as follows: 87 NaCl, 75 sucrose, 25 glucose, 2.5 KCl, 4 MgCl_2_, 0.5 CaCl_2_, 23 NaHCO_3_ and 1.25 NaH_2_PO_4_). Immediately after cutting, slices containing the medial prefrontal cortex (PFC) or the nucleus accumbens were stored for 1 hr at 32°C in a low-calcium ACSF that contained (in mm) as follows: 130 NaCl, 11 glucose, 2.5 KCl, 2.4 MgCl_2_, 1.2 CaCl_2_, 23 NaHCO_3_, 1.2 NaH_2_PO_4_, and were equilibrated with 95% O_2_/5% CO_2_ and then held at room temperature until the time of recording. During the recording, slices were placed in the recording chamber and superfused at 2 ml/min with low Ca^2+^ ACSF. All experiments were done at 32°C. The superfusion medium contained picrotoxin (100 mM) to block GABA-A receptors. All drugs were added at the final concentration to the superfusion medium.

### Electrophysiology

Whole cell patch-clamp of visualized PFC layer five prelimbic pyramidal neurons and field potential recordings were made in coronal slices containing the PFC or the accumbens as previously described (Bara et al. 2018; Borsoi et al. 2019; Scheyer et al. 2019, 2020a, 2020b). Neurons were visualized using an upright microscope with infrared illumination. The intracellular solution was based on K+ gluconate (in mM: 145 K+ gluconate, 3 NaCl, 1 MgCl_2_, 1 EGTA, 0.3 CaCl_2_, 2 Na^2+^ ATP, and 0.3 Na^+^ GTP, 0.2 cAMP, buffered with 10 HEPES). The pH was adjusted to 7.2 and osmolarity to 290 – 300 mOsm. Electrode resistance was 4 – 6 MOhms.

Whole cell patch-clamp recordings were performed with an Axopatch-200B amplifier as previously described (Bara et al. 2018; Borsoi et al. 2019; Scheyer et al. 2019, 2020a, 2020b). Data were low pass filtered at 2kHz, digitized (10 kHz, DigiData 1440A, Axon Instrument), collected using Clampex 10.2 and analyzed using Clampfit 10.2 (all from Molecular Device, Sunnyvale, USA).

Access resistance compensation was not used, and acceptable access resistance was <30 MOhms. The potential reference of the amplifier was adjusted to zero prior to breaking into the cell. Current-voltage (I-V) curves were made by a series of hyperpolarizing to depolarizing current steps immediately after breaking into the cell. Membrane resistance was estimated from the I–V curve around resting membrane potential (Bara et al. 2018; Borsoi et al. 2019; Scheyer et al. 2019, 2020a, 2020b). Field potential recordings were made in coronal slices containing the mPFC as previously described (Bara et al. 2018; Borsoi et al. 2019; Scheyer et al. 2019, 2020a, 2020b). During the recording, slices were placed in the recording chamber and superfused at 2 ml/min with low Ca2+ ACSF. All experiments were done at 32 °C. The superfusion medium contained picrotoxin (100 µM) to block GABA Type A (GABA-A) receptors. All drugs were added at the final concentration to the superfusion medium. The glutamatergic nature of the field EPSP (fEPSP) was systematically confirmed at the end of the experiments using the ionotropic glutamate receptor antagonist CNQX (20 µM), which specifically blocked the synaptic component without altering the non-synaptic component. Both fEPSP area and amplitude were analyzed. Stimulation was performed with a glass electrode filled with ACSF and the stimulus intensity was adjusted ∼60% of maximal intensity after performing an input–output curve. Stimulation frequency was set at 0.1 Hz.

### Data acquisition and analysis

The magnitude of plasticity was calculated at 0-10min and 30–40 min after induction (for TBS-LTP and eCB-LTD) or drug application (mGlu2/3-LTD) as percentage of baseline responses. Statistical analysis of data was performed with Prism (GraphPad Software) using tests indicated in the main text after outlier subtraction (Grubb’s test, alpha level 0.05). All values are given as mean ±SEM, and statistical significance was set at p<0.05.

## Results

To establish the postnatal maturational trajectories layer 5 pyramidal PFC synapses, experiments were performed in rats of both sexes, at the juvenile (female 21<P<28; male 21<P<38), pubescent (female 30<P<50; male 40<P<60) and adult stages (P>90 for both sexes).

### Sex-specific developmental sequence of retrograde endocannabinoid LTD in the rat PFC

Alterations in one or more category of synaptic plasticity exhibited in the PFC have been associated with endophenotypes of neuropsychiatric disorders (Kasanetz et al. 2013; Iafrati et al. 2014; Thomazeau et al. 2014; Labouesse et al. 2017; Manduca et al. 2017). However, most of the above-mentioned studies have been performed solely in adult males. A notable exception is the observation that in-utero cannabinoid exposure selectively ablates endocannabinoid (eCB)-mediated long-term depression (LTD) in the adult male progeny, while females were spared (Bara et al. 2018). Because the PFC and the eCB system both undergo rearrangements during adolescence (Schneider 2008; Bossong and Niesink 2010; Fuhrmann et al. 2015), we elected to systematically investigate eCB-LTD in the PFC of juvenile, pubescent and adult rats of both sexes. First, we confirmed that a 10-minute, 10Hz stimulation of the superficial-layers of the PFC in slices obtained from the adults of both sexes, induces a robust LTD at layer 5 synapses (Figure 1 A-B)(Bara et al. 2018; Borsoi et al. 2019). This identical protocol likewise elicited LTD in slices obtained from pubescent males and females (Figure 1 C-D)(Borsoi et al. 2019). To our surprise, eCB-LTD could not be induced in response to this protocol in the PFC of juvenile males, while juvenile females already expressed full-fledged LTD (Figure 1 E-F).

**Figure 1:**
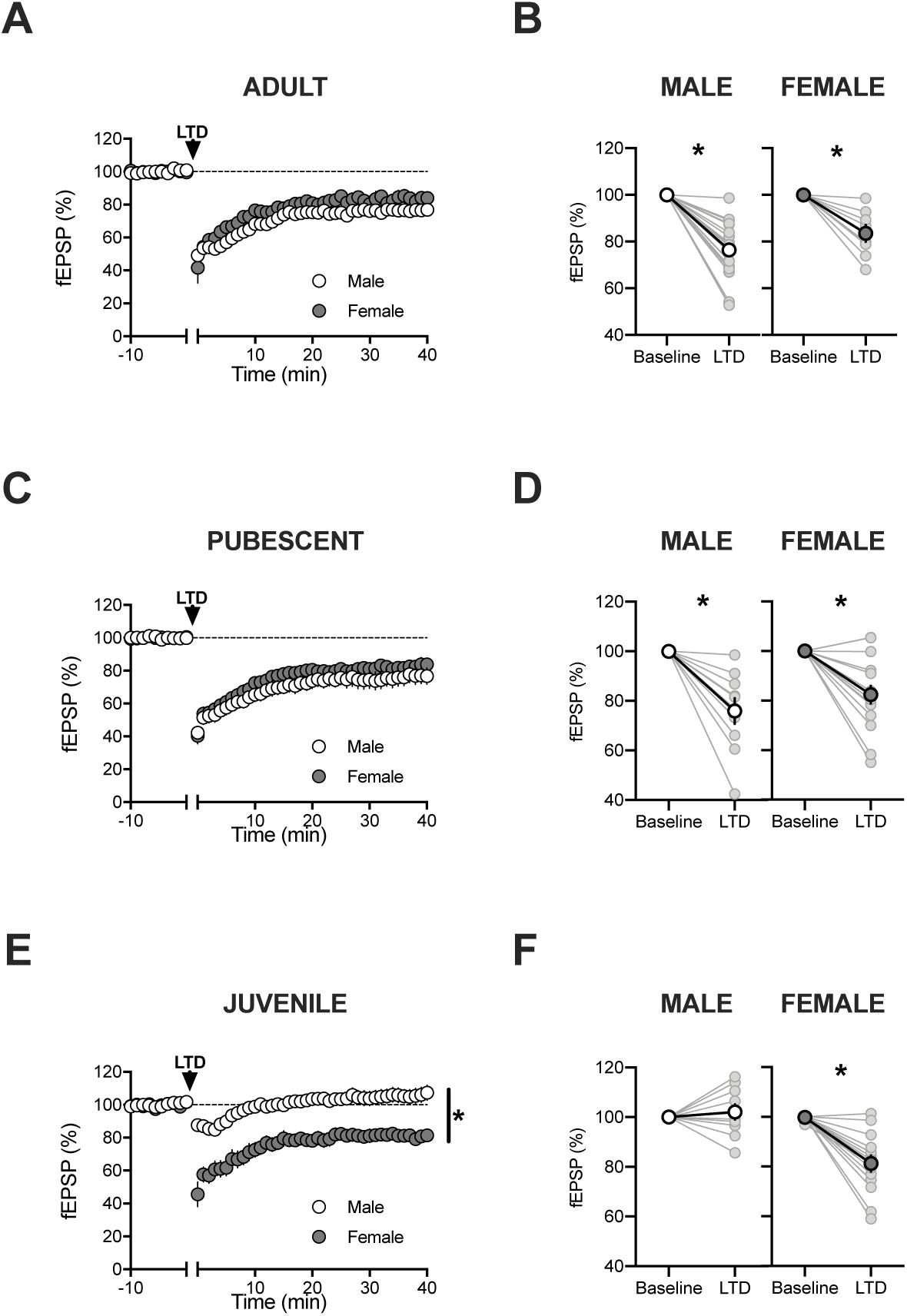
Sex-specific maturational trajectory of endocannabinoid LTD in the rat PFC. **A**. A 10-minute, 10Hz field stimulation (arrow) of layer 2/3 cells in the PFC elicited a robust eCB-LTD at deep layer synapses in adult rat of both sexes. Average time-courses of mean fEPSPs in PFC slices prepared from male (white circles, n=19) or female rats (grey circle, n=8). **B**. fEPSP magnitude at baseline (−10 to 0 minutes) and LTD (30-40 minutes post-tetanus) values corresponding to the normalized values in A. Individual experiments (light circles) and group average (bold circles), before and after LTD induction show similar eCB-LTD in the PFC of adult of both sexes. **C**. Average time-courses of mean fEPSPs showing that a 10-minute, 10Hz field stimulation (arrow) of layer 2/3 cells in the PFC elicited a robust eCB-LTD at deep layer synapses in pubescent rats of both sexes (male, white circles, n=10; female, grey circle, n=16). **D**. Individual experiments (light circles) and group average (bold circles), before and after LTD induction show similar PFC eCB-LTD in pubescent rats of both sexes. **E**. Average time-courses of mean fEPSPs showing the induction of a robust eCB-LTD at deep layer synapses of pubescent female but not male pubescent rats (male, white circles, n=10; female, grey circle, n=14). **F**. Individual experiments (light circles) and group average (bold circles), before and after LTD induction in the juvenile male (left) and female (right) PFC. Data represent mean ± SEM. Two tailed paired T-test, Ordinary one-way ANOVA, Tukey multiple comparisons test, *P<0.05.

Prefronto-accumbens glutamatergic circuits modulate reward-related behaviors (Floresco 2015; Mateo et al. 2017) and the eCB-system of the accumbens is impeded in genetic (Jung et al. 2012) and environmental insults (Lafourcade et al. 2011; Bosch-Bouju et al. 2016) including cannabinoid exposure (Mato et al. 2004, 2005). As shown Figure 2, eCB-LTD was similarly robust in males (Figure 2 A-B) and females at all developmental stages (Figure 2 C-D). Taken together, the results suggest a degree of regional and sex specificity to the maturational profile of eCB-LTD.

**Figure 2:**
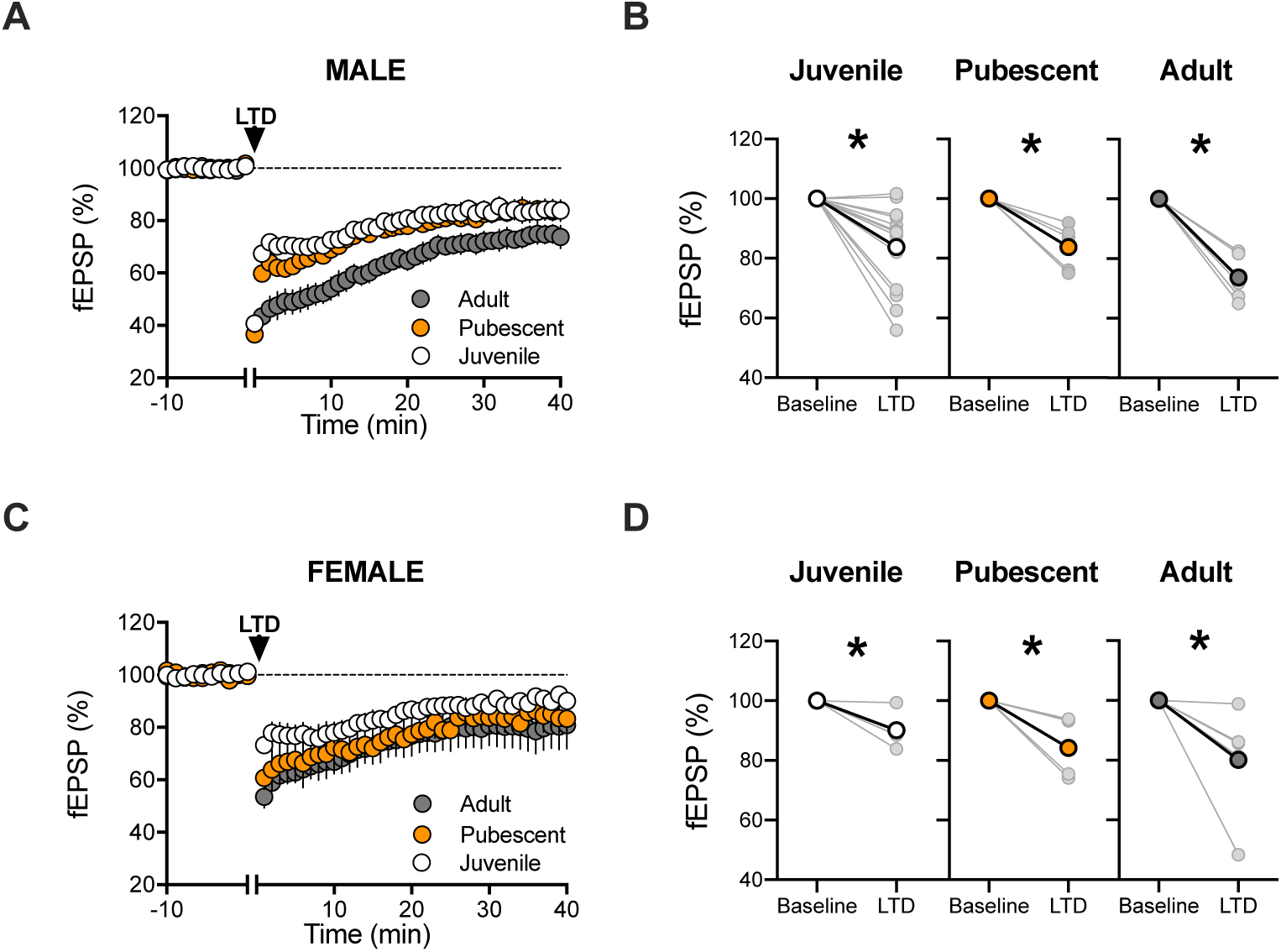
Similar endocannabinoid LTD in the nucleus accumbens core is expressed similarly at all age groups in both sexes. **A**. In male rats, eCB-LTD is induced similarly at accumbens synapses at all age groups in response to the 10-minute, 10Hz field stimulation (arrow) the juvenile to adulthood maturation. Average time-courses of mean fEPSPs in Juveniles (white circles, n=14), Pubescents (orange circles, n=5) and Adults (grey circle, n=5). **B**. Individual experiments (light circles) and group average (bold circles), before (baseline) and after (40min) LTD induction showing similar NAC eCB-LTD in male juvenile, pubescent and adult stages. **C**. In female rats, eCB-LTD is induced similarly at accumbens synapses at all age groups in response to the 10-minute, 10Hz field stimulation (arrow) the juvenile to adulthood maturation. Average time-courses of mean fEPSPs in Juveniles (white circles, n=5), Pubescents (orange circles, n=4) and Adults (grey circle, n=5). **D**. Individual experiments (light circles) and group average (bold circles), before (baseline) and after (40min) LTD induction showing similar NAC eCB-LTD in female juvenile, pubescent and adult stages. Data represent mean ± SEM. Two tailed paired T-test, Two tailed unpaired T-test, Ordinary one-way ANOVA, Tukey multiple comparisons test, *P<0.05.

We next searched for functional alterations that may underlie the delayed expression of eCB-LTD in juvenile males compared to other age groups in males and females.

### CB1R are functional across development and independently of sex

CB1R functionality is sensitive to a number of external factors. For example, nutritional imbalance (Lafourcade et al. 2011) and cannabinoid exposure (Mato et al. 2004, 2005; Mikasova et al. 2008) can desensitize CB1R and consequently ablate eCB-LTD. We tested if functional CB1R were present in both sexes at the three developmental stages by comparing full dose-response curves for the CB1 agonist CP55,940 in both sexes and across age groups. The dose response curves showed that the potency and efficacy of presynaptic CB1R were similar at all ages in both sexes (Figure 3). Thus, the lack of LTD in juvenile males cannot be attributed to a mere lack of presynaptic inhibitory CB1R.

**Figure 3:**
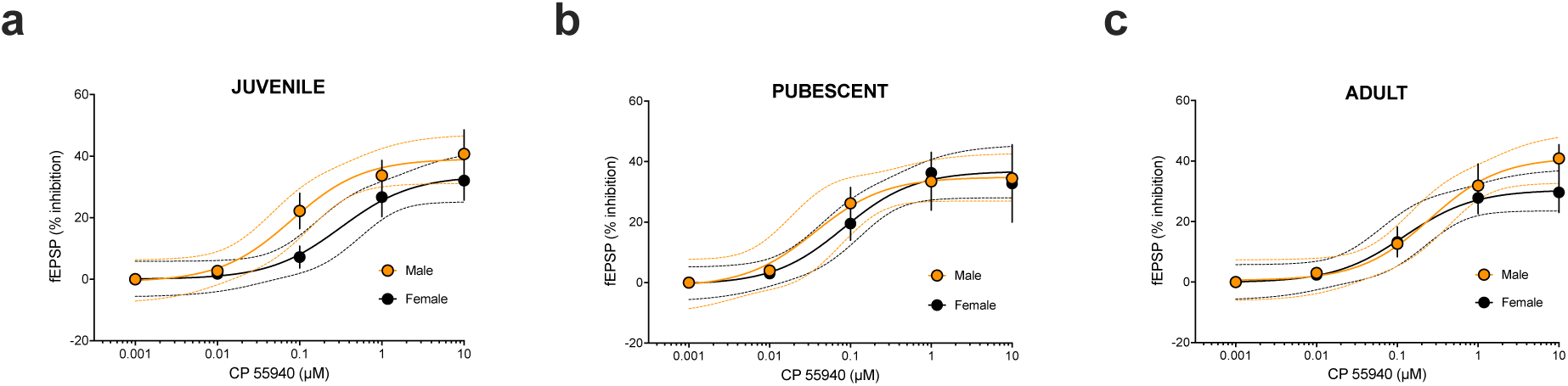
Inhibitory CB1R are similarly functional at all developmental stages in both sexes. **A**. Dose-response curve for the cannabimimetic CP55,940 in juvenile male (orange symbols, n = 3-4 animals, EC_50_ = 0.079 μM, top value 42.17%, 95% CI for EC_50_ = 0.026 – 0.238) and female rats (black symbols, n = 5-6, EC_50_ = 0.316 μM, top value 33.63%, 95% CI for EC_50_ = 0.08 – 1.245). **B**. Dose-response curve for the CP55,940 in pubescent male orange symbols, n = 4-7 animals, EC_50_ = 0.026 μM, top value 34.89%, 95% CI for EC_50_ = 0.006 – 0.204) and female rats (black symbols, n = 4-6, EC_50_ = 0.082 μM, top value 36.85%, 95% CI for EC_50_ = 0.027 – 0.255). **C**. Dose-response curve for the CP55,940 in adult male (orange symbols, n = 3-5 animals, EC_50_ = 0.253 μM, top value 41.21%, 95% CI for EC_50_ = 0.0813 – 0.7853) and female rats (black symbols, n = 5-6, EC_50_ = 0.122 μM, top value 30.5%, 95% CI for EC_50_ = 0.0296 – 0.504). fEPSP amplitudes were measured 30 min after application of CP55,940. Each point is expressed as the percentage of inhibition of its basal value. Error bars indicate SEM.

### Sex-specific maturation of PFC pyramidal cell excitability and basic synaptic properties

In search of mechanistic insights, we compared intrinsic firing properties of pyramidal neurons in our various experimental groups. We performed patch-clamp recordings of deep-layer PFC pyramidal neurons in acute PFC slices obtained from juvenile, pubescent and adult male and female rats and measured the membrane reaction profiles in response to a series of somatic current steps. Independent of developmental stage or sex all recorded layer 5 PFC neurons showed similar and superimposable *I– V* plots (Fig. 4 A-B). Similarly, the resting membrane potential (Fig. 4 C-D) and the rheobase (Fig. 4 E-F) were alike within and between age and sex groups. Closer examination of the depolarization-driven action potentials showed less action potentials in response to somatic current steps, in female adults compared to the other developmental stages (Figure 4 G-H). This trait, while suggestive of a lower excitability of PFC pyramidal neurons in adult females, does not provide a mechanism for the lack of LTD in juvenile male.

**Figure 4:**
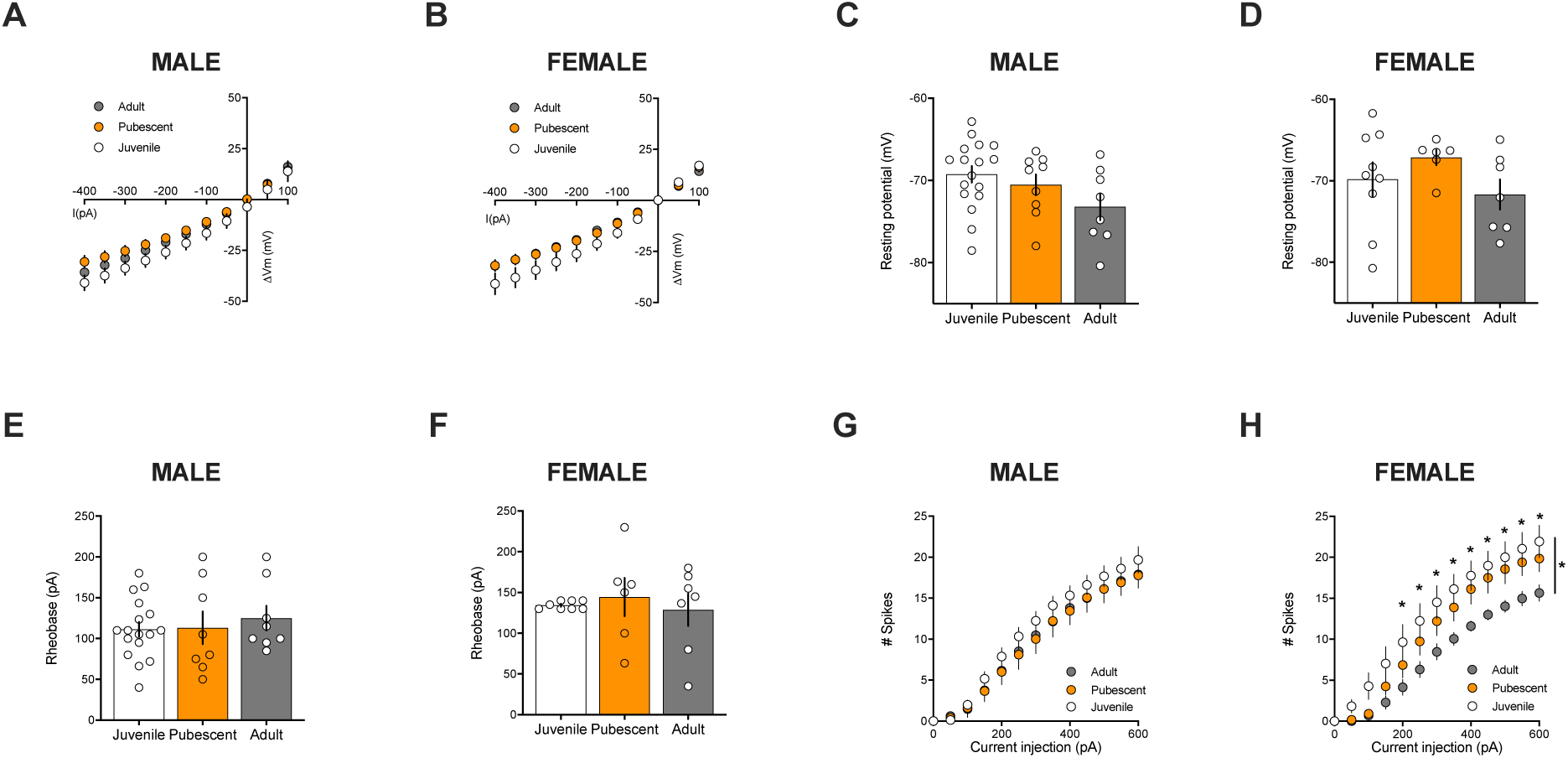
Sex-specific maturation of the intrinsic properties of layer 5 PFC pyramidal neurons. **A-B**. Comparison of current injection steps of 50pA from -400pA to 100pA revealed no differences in the I-V relationship in layer 5 pyramidal neurons of the PFC of female and male groups at the juvenile, pubescent and adult stages. **C-D**. Similarly, no difference was found in resting membrane potentials and **(E-F)** rheobases (i.e. the minimum current injection required to elicit an action potential with 10pA progressive current injections steps from 0-200pA). **G**. In male, the number of evoked action potentials in response to increasing depolarizing current steps from 0-600pA did not differ in PFC pyramidal neurons of male rats during the juvenile to adulthood maturation. **H**. In female however, the number of evoked action potentials in response to increasing depolarizing current steps was lower in adults versus juvenile and pubescent. Data represent mean ± SEM. Scatter dot plot represents one animal. Males (Juvenile n=16; Pubescent n=9; Adult n=9); Females (Juvenile n=9; Pubescent n=6; Adult n=7). n= individual animal. Sidak’s multiple comparisons test, *P<0.05.

Basic synaptic properties were also compared. Input–output profiles were measured in the 6 groups: fEPSPs evoked by electrical stimulation showed a consistent profile distribution in response to increasing stimulation intensity: “input–output” curves were similar across age and sex (Fig. 5 A-C). Likewise, the paired-pulse ratio (PPR), a multifactorial parameter of the presynaptic probability of neurotransmitter release, remained unchanged in the experimental groups. When testing whether the PPR changed throughout development, we observed that paired stimuli elicited with intervals >150 ms induced equivalent facilitation in juveniles, pubescent and adults (Figure 5 D-E). Thus, the release probability of excitatory synapses to PFC layer 5 pyramidal neurons, as estimated from the PPR, was stable throughout developmental stages and in both sexes.

**Figure 5:**
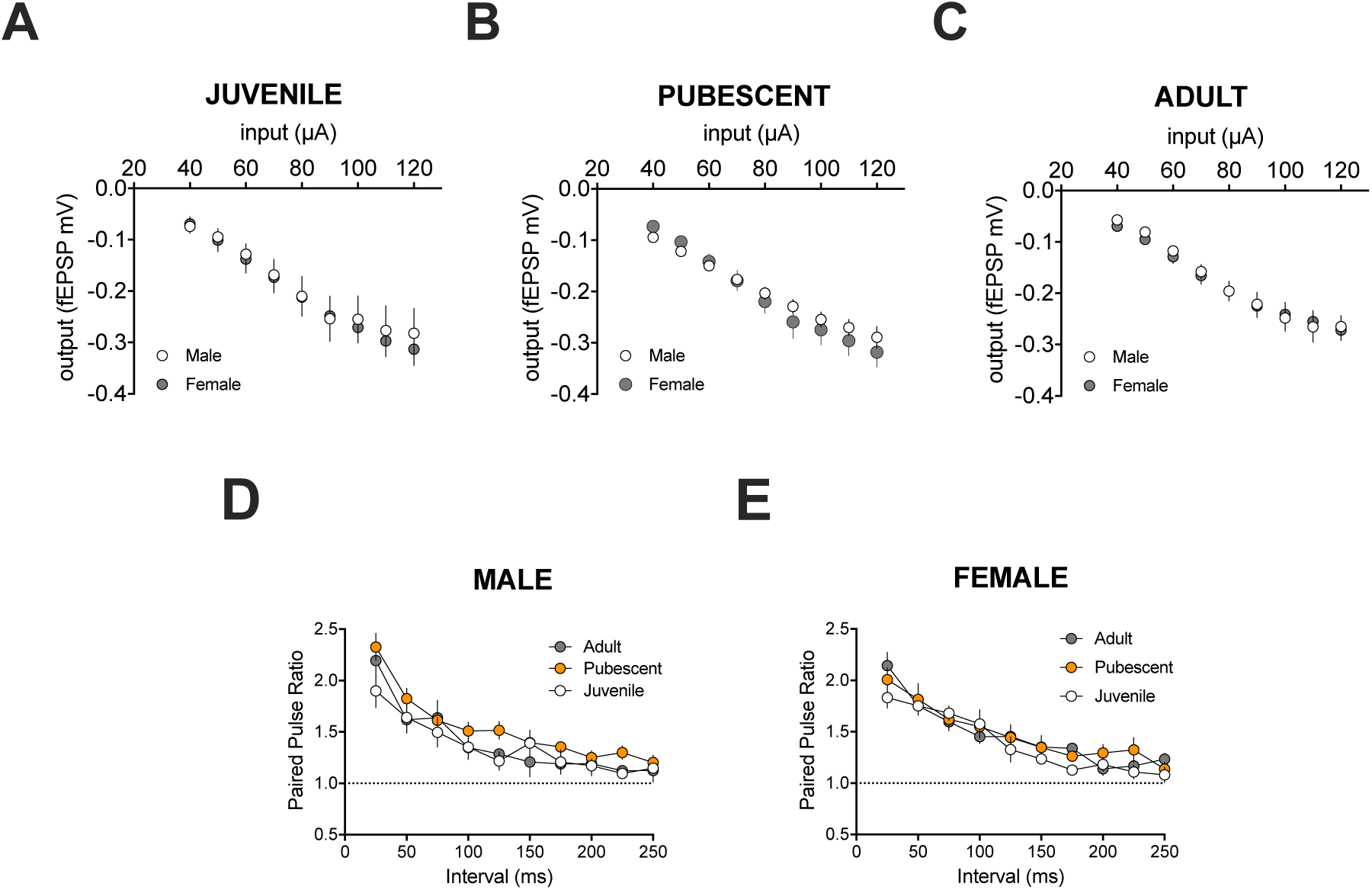
The basic synaptic properties of mPFC pyramidal are identical at all age groups in both sexes. **A-C**. Input–output profile from juvenile (A), pubescent (B) and adult mPFC layer 5 fEPSPs. Averaged fEPSP area measured as a factor of stimulus intensity. **D-E**: Short-term plasticity of fEPSPs estimated by the ratio of paired stimulus-induced fEPSPs normalized to the amplitude of the first response in juvenile, pubescent and adult male (**D**) or female (**E**) rats showed no modification in PPR ratio curves along late postnatal development. n= individual animal. Sidak’s multiple comparisons test.

### Sexually dimorphic mechanisms of LTD

In an earlier study, we revealed a previously unknown sexual difference in the mechanism of PFC eCB-LTD: LTD is mediated solely by CB1R in males, in striking contrast to females where activation of TRPV1R (but not CB1R) is required to elicit LTD (Bara et al. 2018). We reproduced and extended this observation to show that eCB-LTD engages distinct receptors in male and female depending on their developmental stages (Figure 6). In pubescent and adult males, the induction of LTD was blocked by the CB1R antagonist (SR141716A, Figure 6 A-B) but not by the TRPV1R antagonist (AMG 9810, Figure 6 C-D). Thus, regardless of the developmental stage, males’ eCB-LTD solely depends on CB1R. In marked contrast, either AMG9810 or SR141716A could prevent LTD induction in juvenile and pubescent females (Figure 6 E-H). Female pubescent LTD was only sensitive to AMG9810 (Figure 6G-H). As shown before (Bara et al. 2018), LTD was unaffected by the CB1R antagonist but blocked by AMG9810 in adult females (Figure 6 G-H). Thus, female rats require both CB1R and TRPV1R to make eCB-LTD as juveniles, only CB1R during the pubescent stage, and then solely TRPV1R at adulthood.

**Figure 6:**
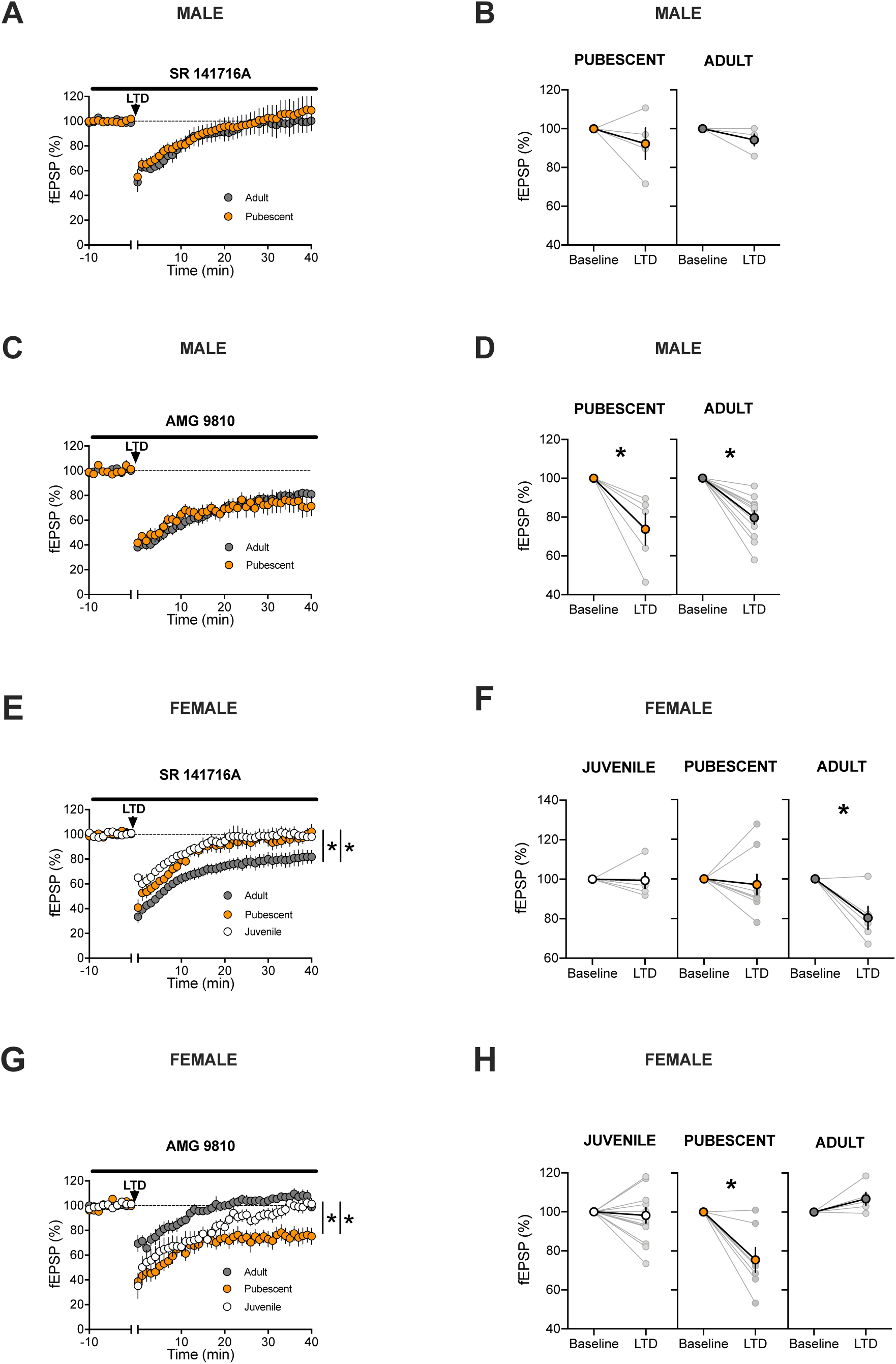
Sex specific mechanisms of LTD across postnatal development. **A-D**. In males’ PFC, LTD is always mediated by CB1R. **A**. In pubescent and adult males, a > 45 min pre-incubation with the CB1R antagonist (SR141716A) prevents the induction of 10Hz LTD (arrow). Average time-courses of mean fEPSPs in PFC slices prepared from SR141716A-treated pubescent (orange circles, n=5) and adult (grey circle, n=5) males. **B**. fEPSP magnitude at baseline (−10 to 0 min) and LTD (30-40 min post-tetanus) values. Individual experiments (light circles) and group average (bold circles) in the PFC of male pubescent and adult rats. **C**. In pubescent and adult males, a > 45 min pre-incubation with the TRPV1R antagonist (AMG9810) had no effect on LTD induction (arrow). Average time-courses of mean fEPSPs in PFC slices prepared from AMG9810 pubescent (orange circles, n=5) and adult (grey circle, n=11) males. **D**. fEPSP magnitude at baseline and LTD values. Individual experiments (light circles) and group average (bold circles) in the PFC of pubescent and adult male rats. **E-H**. In females’ PFC, back-and-forth CB1R and TRPV1R are required for LTD **E**. In juvenile and pubescent female, the CB1R antagonist prevents LTD induction, while the same protocol elicited a robust eCB-LTD in adult females. Average time-courses of mean fEPSPs in PFC slices prepared from SR141716A-treated juvenile (white circles, n=5) pubescent (orange circles, n=9) and adult (grey circle, n=6) females. **F**. fEPSP magnitude at baseline and LTD. Individual experiments (light circles) and group average (bold circles) after SR141716A in the PFC of pubescent and adult females. **G**. In juvenile and adult but not pubescent females, the TRPV1R antagonist prevented eCB-LTD. Average time-courses of mean fEPSPs in PFC slices prepared from AMG9810-tretaed juvenile (white circles, n=13) pubescent (orange circles, n=7) and adult (grey circle, n=5) females **H**. fEPSP magnitude at baseline and LTD. Individual experiments (light circles) and group average (bold circles) show eCB-LTD after AMG9810 incubation in the PFC of female juvenile, pubescent and adult rats. Data represent mean ± SEM. Two tailed paired T-test, Two tailed unpaired T-test, Ordinary one-way ANOVA, Tukey multiple comparisons test, *P<0.05.

### Blocking the degradation of 2AG uncovers LTD in juvenile male PFC

In the rodent PFC, eCB-LTD requires the participation of 2AG and/or anandamide (present data, Bara et al. 2018). In contrast to anandamide that activates both TRPV1R and CB1R, 2AG selectively stimulates CB1R. The current data showing that CB1R mediates LTD in pubescent and adult male rats, open the possibility that enhancing 2AG levels could allow for the induction of eCB-LTD in juvenile males. Thus, PFC slices were incubated (>45 min) in JZL184 (4 µM) a potent inhibitor of monoacyl-glycerol lipase, the main enzyme degrading 2AG, in order to increase basal 2-AG levels prior to the 10-minute, 10Hz stimulation. Here, after JZL184 incubation slices obtained from juvenile males were found to exhibit robust, lasting LTD at layer 5 synapses (Figure 7). Thus, increasing the levels of 2-AG in the PFC effectively uncovers eCB-LTD in the otherwise immature male system.

**Figure 7:**
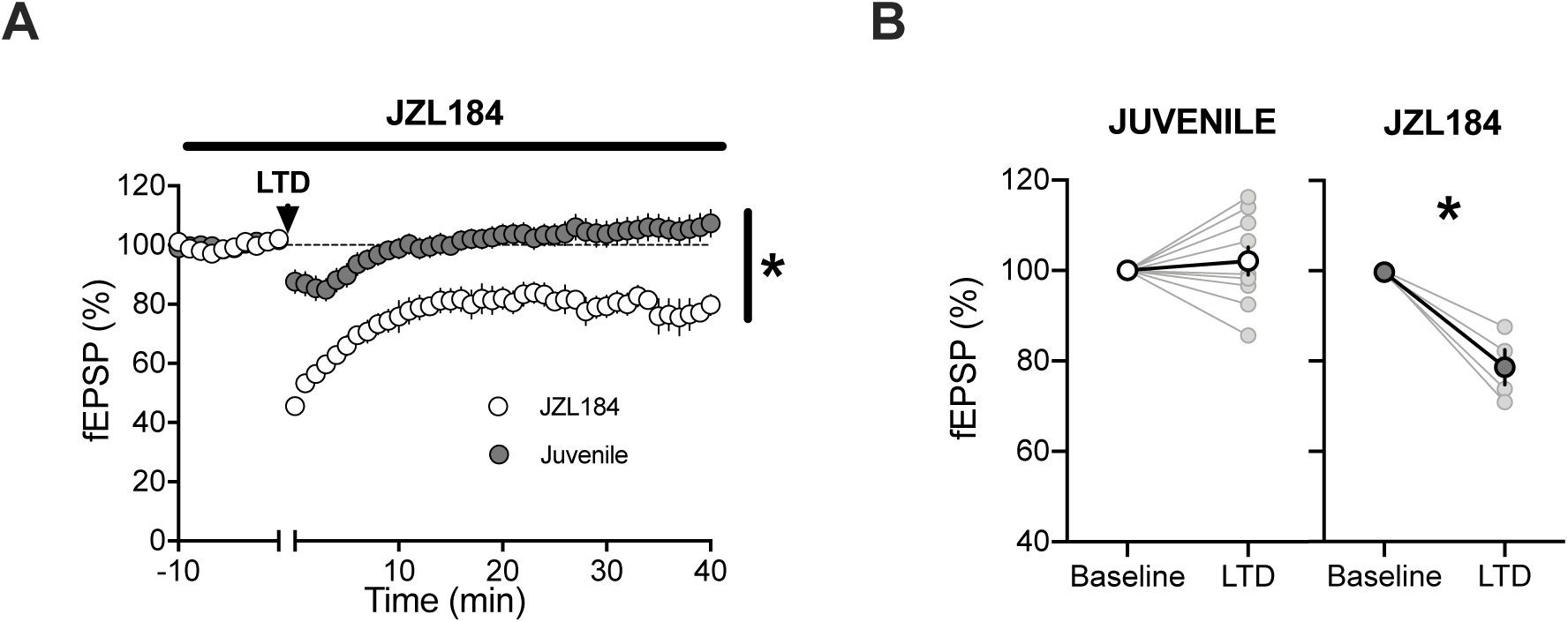
Enhancing 2-AG with JZL184 effectively uncovers eCB-LTD in the juvenile male PFC. **A**. A 10Hz field stimulation (arrow) of layer 2/3 cells in the PFC of male juvenile rats failed to induce eCB-LTD at deep layer synapses. However, following a > 45 min pre-incubation with the MAGL inhibitor, JZL 184 (1µM), the previously ineffective low frequency protocol now induced a robust LTD in male juvenile rats. Average time-courses of mean fEPSPs in PFC slices prepared from JZL 184 treated (white circles, n=4) or untreated (grey circle, n=10) juvenile males. **B**. fEPSP magnitude at baseline (−10 to 0 minutes) and LTD (30-40 minutes post-tetanus) values corresponding to the normalized values in A. Individual experiments (light circles) and group average (bold circles) show eCB-LTD after JZL 184-incubation in the PFC of male juvenile rats. Data represent mean ± SEM. Two tailed paired T-test, Two tailed unpaired T-test, *P<0.05.

### Long-term potentiation and mGluR2/3 LTD do not follow a sex-specific maturational sequence in the rat PFC

We next determined if the sex-specific maturation of LTD in the PFC was global or limited to eCB-LTD. We examined a distinct form of LTD in the rat PFC mediated by mGlu2/3 receptors (Otani et al. 2002; Huang et al. 2007; Kasanetz et al. 2013; Borsoi et al. 2019). mGlu2/3 LTD and eCB-LTD that share common presynaptic mechanisms (Mato et al. 2005) and thus may display similar sex-specific maturational paths. PFC slices from our various groups were exposed for 10-minutes to the mGlu2/3 agonist, LY379268 (300nM) in order to induce an mGlu2/3-dependent LTD. As shown before (Borsoi et al. 2019), this drug application elicited a significant LTD at layer 5 synapses in slices obtained from adult (Figure 8 A-B) or pubescent (Figure 8 C-D) rats of both sexes (Figure 8 A-B). Similarly, the 10-minute application effectively elicited LTD in slices obtained from juvenile male and female rats (Figure 8 E-F). Thus, in spite of shared downstream pathways, mGlu2/3-LTD does, not follow the sex-specific maturational trajectory of eCB-LTD in the PFC of rats.

**Figure 8:**
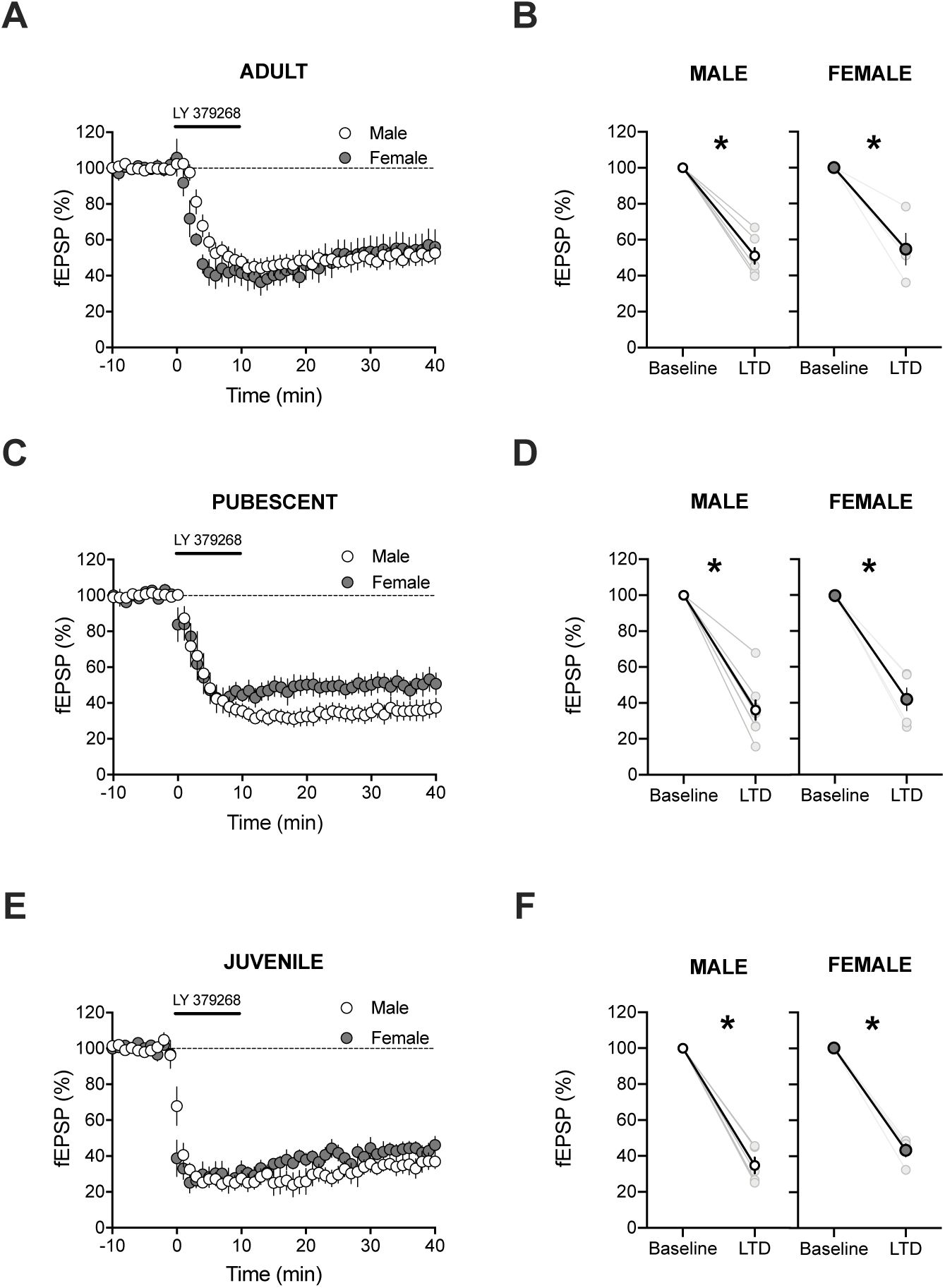
mGluR2/3-LTD does not follow a sex-specific sequence in the rat PFC. **A**. mGluR-LTD, induced by a 10-min application of 30nM LY379268, produced a significant LTD at deep layer synapses in adult rat of both sexes. Average time-courses of mean fEPSPs in PFC slices prepared from male (white circles, n=6) or female rats (grey circle, n=4). **B**. fEPSP magnitude at baseline (−10 to 0 minutes) and LTD (30-40 minutes post-drug washout) values corresponding to the normalized values in A. Individual experiments (light circles) and group average (bold circles), show similar mGluR2/3-LTD in the PFC of adult of both sexes. **C**. Average time-courses showing mGluR2/3-LTD in the PFC of pubescent rats of both sexes (male, white circles, n=8; female, grey circle, n=5). **D**. Individual experiments (light circles) and group average (bold circles), before and after mGluR2/3-LTD induction in pubescent rats of both sexes. **E**. Average time-courses showing the induction of a robust mGluR2/3-LTD in pubescent male and female rats (male, white circles, n=5; female, grey circle, n=4). **F**. Individual experiments (light circles) and group average (bold circles), before and after mGluR2/3-LTD induction in the juvenile male (left) and female (right) PFC. Data represent mean ± SEM. Two tailed paired T-test, Two tailed unpaired T-test, Ordinary one-way ANOVA, Tukey multiple comparisons test, *P<0.05.

Next, we elected to investigate a third form of plasticity in the PFC of both sexes at the three age periods in order to determine the extent of the sexual dimorphism of PFC plasticity. We evaluated a plasticity known to be altered in the PFC of rodent models of neuropsychiatric disorders (Thomazeau et al. 2014; Iafrati et al. 2016; Labouesse et al. 2017; Manduca et al. 2017), the NMDAR-dependent theta-burst induced long-term potentiation (TBS-LTP). A TBS was applied to superficial layers of acute PFC slices obtained from our various rat groups, during simultaneous recording of extracellular field EPSPs at deep layer synapses. The TBS protocol effectively induced a lasting synaptic potentiation in slices obtained from rats of all age periods and both sexes (Figure 9), thereby confirming and extending a previous study (Borsoi et al. 2019). The data indicate that TBS-LTP, is mature at the juvenile stage in rats of both sexes and consistently expressed in pubescent and adult of both sexes.

**Figure 9:**
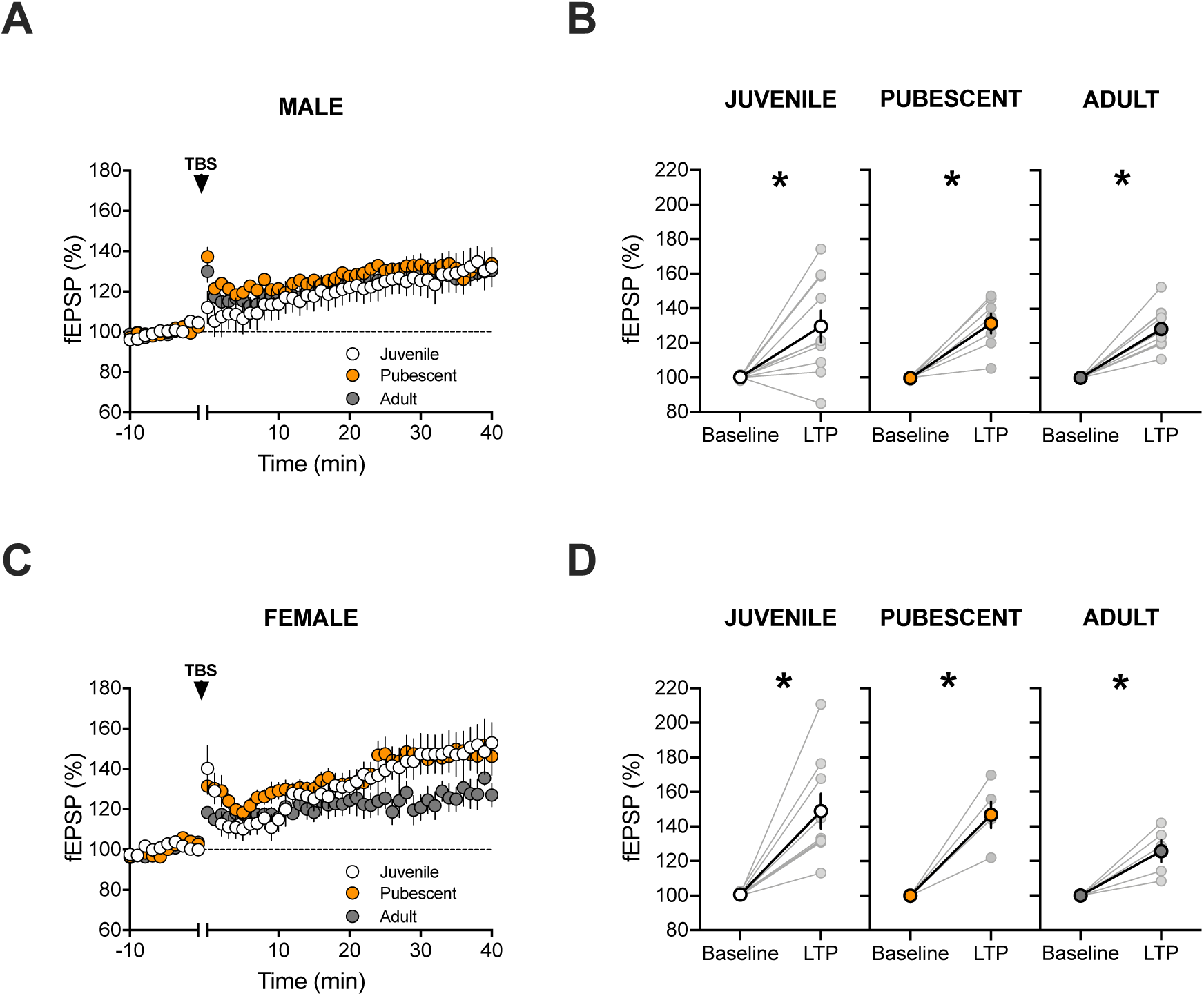
LTP does not follow a sex-specific sequence in the rat PFC. **A**. In male PFC, theta-burst stimulation (arrow, 5 pulses @ 100 hz, 4 times) of layer 2/3 cells in elicited a robust LTP at deep-layer synapses at all age groups. Average time-courses of mean fEPSPs in PFC slices prepared from juvenile (white circles, n=10), pubescent (orange circle, n=7) and adult (grey circle, n=11) male rats. **B**. Individual experiments (light circles) and group average (bold circles), before and after LTP induction show similar LTP at all stages. **C**. In female rats, LTP is induced similarly in juvenile, pubescent and adult groups. Average time-courses of mean fEPSPs in Juveniles (white circles, n=9), Pubescents (orange circles, n=7) and Adults (grey circle, n=5). **D**. Individual experiments (light circles) and group average (bold circles) show similar LTP in female at all stages. Data represent mean ± SEM. Two tailed paired T-test, Two tailed unpaired T-test, Ordinary one-way ANOVA, Tukey multiple comparisons test, *P<0.05.

## Discussion

We applied a cross-sectional strategy to show that in deep-layer (L5) pyramidal neurons of the rat PFC, the maturation of excitatory input endocannabinoid-mediated synaptic plasticity follows a sex-specific trajectory throughout the juvenile to adult stages. Specifically, we show that while eCB-mediated LTD is already apparent in juvenile females, it is expressed after the onset of puberty in males. The age-related and sex-specific regulation of eCB plasticity contrasts with other forms of synaptic plasticity examined that were already mature by the time of birth in both sexes.

By when examining their postnatal trajectories, we observed relatively few changes in the intrinsic properties of PFC principal neurons. Features such as resting membrane potentials and rheobase did not vary across age and sex, with the exception that the number of action potentials in response to depolarizing steps was reduced in adult females compared to all other groups, suggesting decreased excitability in the adult females. Similarly to intrinsic membrane properties, two basic synaptic parameters, the paired-pulse ratio and the input-output relationships, were remarkably stable across maturation and sexes. These results suggest that for the most part, cellular and synaptic properties of PFC neurons are established early in life and that the inability to induce eCB-LTD in juvenile males (see below) cannot be attributed to the above-mentioned characters.

Early life and adolescence are well described periods of changes for the eCB system (Ellgren et al. 2008; Heng et al. 2011; Lee and Gorzalka 2012; Rubino and Parolaro 2015; Meyer et al. 2018). The current data support and extend this concept by showing that a major form synaptic plasticity mediated by eCBs is also subject to developmental regulation and is the substrate of a previously unrecognized sex difference before puberty. Furthermore, the observation that eCB-LTD was consistently expressed in both sexes and at all developmental stages in the accumbens strongly suggests that this sexual dimorphism is specific to the PFC. The PFC achieves maturity late into adolesce and additional work will determine the role of eCB-LTD in the behavioral attributes of adolescence.

Dysfunctional CB1R could underlie the lack of eCB-LTD in juvenile male. Indeed, in their pioneering studies, Rodriguez de Fonseca and colleagues observed sex differences in CB1R expression starting at early life and peaking around adolescence (Rodríguez de Fonseca et al. 1993, 1994). We evaluated the functionality of presynaptic CB1R by building dose–response curves for the cannabinoid agonist CP55,940. Maximal inhibitory effects of CP55,940 and its EC50 were comparable in all our experimental groups (Fig. 3) and similar to what we previously reported in adult male rats (Bara et al., 2018). Thus, even though CB1R expression peaks with the onset of adolescence (Meyer et al. 2018), one can reasonably exclude the possibility that the lack of PFC eCB-LTD in juvenile males is due to insufficient or non-functional presynaptic CB1R.

Circulating levels of anandamide and 2-AG fluctuate and peak around adolescence (Ellgren et al. 2008; Heng et al. 2011; Lee and Gorzalka 2012; Rubino and Parolaro 2015; Meyer et al. 2018). In this context, the expression of eCB-LTD following MAGL inhibition in juvenile males and the expected enhanced 2-AG levels extends and confirms previous similar findings in other models of genetic diseases or environmental insults (Jung et al. 2012; Thomazeau et al. 2014; Bosch-Bouju et al. 2016). This result is also in line with the reported developmental profile of MAGL (Meyer et al. 2018).

Cannabis exposure during adolescence produces multiple long-lasting and sex-specific molecular, synaptic and behavioral alterations (Cass et al. 2014; Rubino and Parolaro 2015; Renard et al. 2016; Borsoi et al. 2019). Based on the current findings it is tempting to attribute at least partially, such differences to underlying sex and state-dependent differences in the maturational window of development during which cannabinoid exposure takes place.

Additional behavioral and molecular investigations as well as extended characterizations of plasticity and synaptic functions in animals of both sexes in other brain regions are necessary to provide a more thorough picture of the extent to which adolescent maturation shapes brain functions in adulthood.

## Author contributions

Axel Bernabeu: Conceptualization, Data curation, Formal analysis, Validation, Writing—review and editing.

Anissa Bara: Conceptualization, Data curation, Formal analysis, Validation, Writing— review and editing.

Antonia Manduca: Data curation, Formal analysis, Validation.

Milene Borsoi: Data curation, Formal analysis, Validation.

Olivier Lassale: Data curation, Formal analysis, Validation, Methodology.

Anne-Laure Pelissier-Alicot: Conceptualization, Supervision, Funding acquisition, Project administration, Editing.

Olivier JJ Manzoni: Conceptualization, Supervision, Funding acquisition, Methodology, Writing—original draft, Project administration, Writing—review and editing.

The authors declare no conflict of interest.

## Funding and Disclosures

This work was supported by the Institut National de la Santé et de la Recherche Médicale (INSERM); Fondation pour la Recherche Médicale (Equipe FRM 2015 to O.M.) and the NIH (R01DA043982 to O.M.).

## Declarations of interest

The authors declare no competing interests.

## Acknowledgements

The authors are grateful to the Chavis-Manzoni team members for helpful discussions and to Drs. A.F. Scheyer and K. Mackie for critical reading and help with writing the manuscript.

## Notes

### Competing Interest Statement

The authors have declared no competing interest.

### Summary of Updates

The main text was updated and corrected.

